# Most protein domains exist as variants with distinct functions across cells, tissues, and diseases

**DOI:** 10.1101/2022.08.12.503740

**Authors:** Kristoffer Vitting-Seerup

## Abstract

**Background:** Protein domains are the active subunits that provide proteins with specific functions through precise three-dimensional structures. Such domains facilitate most protein functions, including molecular interactions and signal transduction. Currently, these protein domains are described and analyzed as invariable molecular building blocks with fixed functions.

**Results:** I show that most human protein domains exist as multiple distinct variants that I term “domain isotypes”. Different domain isotypes are used in a cell, tissue, and disease-specific manner and have surprisingly different 3D structures. Accordingly, I find that domain isotypes, compared to each other, modulate or abolish the functionality of protein domains.

**Conclusions:** My results challenge the current view of protein domains as invariable building blocks and have significant implications for both wet- and dry-lab workflows. The extensive use of protein domain isotypes within protein isoforms adds to the growing body of evidence suggesting that the sciences should move from the current gene-centric research paradigm toward an isoform-centric research paradigm.

## Introduction

Most proteins have, throughout evolution, been created using protein domains as molecular building blocks (Aziz and Caetano-Anollés, 2021). The importance of these protein domains is hard to overstate as they encode the core functionality needed for most if not all, cellular functions. Noteworthy examples include signal transduction (Roskoski, 2004) and the ability of proteins to bind to DNA, RNA, and other proteins (Mosca et al., 2014). Current state-of-the-art annotation of protein domains (e.g., Pfam (Mistry et al., 2020)) relies on encapsulating known domain sequences into a reference model and matching those models to a sequence of interest. Domain annotation tools are used in many types of analysis and range from mechanistic studies of proteins to interpreting the impact of mutations and are cited by thousands of papers each year.

While protein domains are defined, described, and analyzed as protein subunits with fixed functionality (Jones et al., 2014; Mistry et al., 2020), that might not be the case. Numerous mutational and cancer studies find that removing just a tiny part of a protein domain can either eliminate or modify domain function (Kato et al., 2003; Klimovich et al., 2022; Oren and Rotter, 2010). Such findings lead to an intriguing hypothesis: just like most genes produce protein isoforms with distinct functions, naturally occurring protein domain variants could exist. Such domain variants would originate from alternative splicing or evolution, resulting in different isoforms or genes with variants of the same protein domain.

Here I term this phenomenon “domain isotypes” and show that in humans, they are ubiquitous, used in a cell, tissue, and disease-specific manner, have distinct 3D structures, and can modify the biological function of a protein domain.

## Results

### Domain isotypes exist in humans

To investigate if alternative splicing creates domain isotypes in humans, I first analyzed dysregulated splicing in human cancers. Specifically, I focused on cases where tumors, compared to adjacent healthy tissue, had upregulated one isoform while downregulated another (often referred to as isoform switches) (Vitting-Seerup and Sandelin, 2017). I re-analyzed these isoform switches for cases where the difference between the isoforms was a partial domain loss/gain. Across the 12 cancer types analyzed, I identified 492 genes where an isoform switch resulted in substantial changes to, but not complete gain/removal of, the protein domain (Figure S1A-1, Table S1). Notably, 204 of these genes were found in more than one cancer type (Figure S1A-2), highlighting their functional importance.

A prominent example is the isoform switch in the STK4 gene (frequently called MST1) found in 3 cancer types: thyroid, kidney, and colorectal (Figure S1A-3, S1A-4, S1A-5). STK4 is a key component in the Hippo pathway (Thompson and Sahai, 2015), which functions as a tumor suppressor by phosphorylating and inactivating the YAP1 oncogene (Patel et al., 2017). Inactivation of STK4 has been linked to poor patient prognosis in many cancer types, including colon and kidney cancer (Cinar et al., 2021; Han, 2019; Rybarczyk et al., 2017). The isoform switch I identified in STK4 results in the central part of the Pkinase domain, including the active sites, being removed from the cancer isoform (Figure 1A, S1A-3). Since the Pkinase domain is responsible for the phosphorylation of YAP1, the cancer isoform most likely cannot inhibit YAP1, potentially leading to cancer progression, and suggests domain isotypes, created via alternative splicing, are important in human cancers.

**Figure 1:**
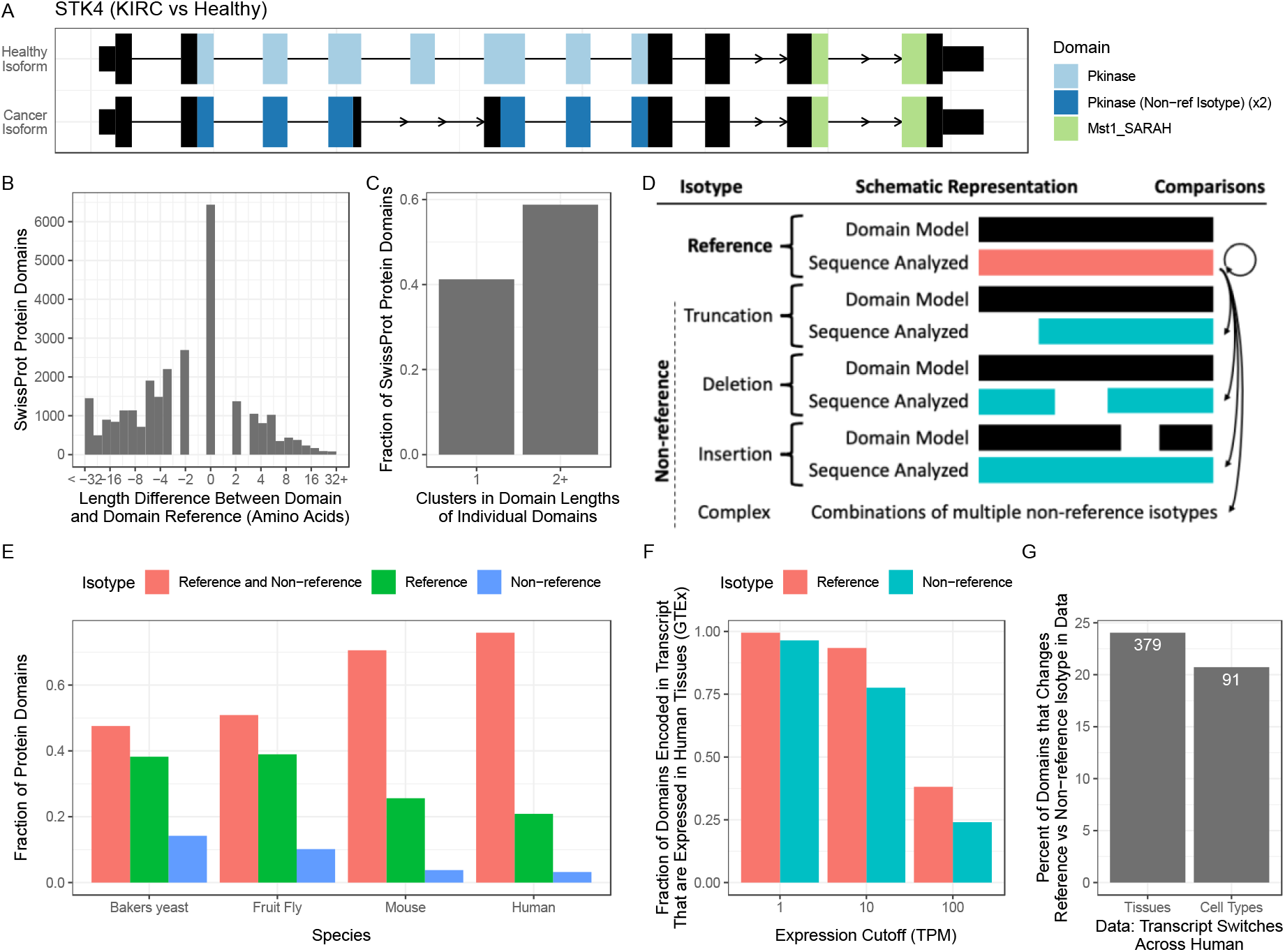
Domain isotypes are ubiquitous. **A)** The isoform switch in ABI1 identified by comparing Lung Squamous Cell Carcinoma (LUSC) to adjacent healthy tissue. Boxes represent exons; where the higher sections annotate, the coding regions and protein domains are indicated by color. The isoform usage is indicated on the left side. Note the differences in the Abi_HRR domain. **B)** The difference between the actual length of protein domains and their corresponding reference length across all human domains identified in SwissProt. **C)** The number of clusters found in the length distribution of each human protein domain. **D)** Schematic representation of the five different protein domain isotypes identified. Black boxes represent the reference model, and red/turquoise boxes indicate the sequence analyzed (red being the reference isotype and turquoise being non-reference isotypes). Arrows indicate comparisons analyzed in this paper; the non-reference isotypes are highlighted to the left. **E)** The fraction of protein domains that exist as reference and non-reference isotypes in SwissProt of various species. **F)** The fraction (y-axis) of protein domain isotypes (color) contained in transcripts expressed above a cutoff (x-axis) in the average expression profile of at least one human tissue. **G)** The most expressed transcript was identified from each gene in each human tissue, and genes where the major transcript differed across tissues or cell types (x-axis) were extracted. From these transcripts, I calculated the percent of encoded protein domains (Y-axis) where there was a domain isotype switch due to the transcript switch. Numbers on the bars indicate the actual number of domains identified.

Encouraged by these examples, I next asked if domain isotypes are also created through evolution. I, therefore, analyzed the length of all protein domains identified in the Swiss-Prot canonical database (11076 domain-contain proteins, one isoform per gene)(UniProt-Consortium et al., 2020). Compared to the domain length annotated in the Pfam database (Mistry et al., 2020), I found substantial length differences in the domains identified in humans (Figure 1B). Intriguingly many protein domains seem to follow individual patterns in how they deviate from the reference length (Figure S1C-1). To investigate this, I used the fact that most protein domains are found many times in the human proteome and extracted the domain length for each of these observations. For each protein domain, I then used cluster analysis to look for patterns in the length distribution. For most protein domains (58.3%), the length of the domains separates into two or more clusters (Figure 1C, S1C-2), suggesting domain isotypes are frequently created and used through evolution.

In summary, I find that alternative splicing an evolution creates domain isotypes.

### Enabling systematic analysis of domain isotypes

To systematically analyze domain isotypes, I developed pfamAnalyzeR, which enables detection and in-depth analysis of domain isotypes (available via Bioconductor (Huber et al., 2015) at https://bioconductor.org/packages/pfamAnalyzeR/). pfamAnalyzeR uses a highly stringent approach to detect domain changes (Figure S1D-1) and can distinguish between 5 categories of domain isotypes (Figure 1D). These isotypes are the reference isotype and four isotypes that, compared to the reference isotype, are best described as a truncation, an insertion, a deletion, or combinations thereof (“complex”)(See schematic illustration in Figure 1D). Importantly the stringent cutoff used here also means that my results cannot be explained by erroneous Pfam domain identification such as those identified by Triant *et al*. (Triant and Pearson, 2015) as those, if truly erroneous, represent less than 0.1% of the data I here analyze.

### Domain isotypes are ubiquitous

I used pfamAnalyzeR to analyze all protein domains in the manually curated Swiss-Prot database of multiple species (UniProt-Consortium et al., 2020). Intriguingly I find that most protein domains, as a consequence of evolution, exist as both reference and non-reference isotypes (Figure 1E, S1E-1, Table S2). Importantly these frequencies are even higher among the proteins annotated as “domain containing” by the Swiss-Prot curators suggesting these are not due to erroneous annotation (Figure S1E-2).

### Domain isotypes are used in a cell and tissue-specific manner

To assess the biological relevance of domain isotypes, I examined the expression of mRNA transcripts containing either reference or non-reference domain isotypes across the 55 human tissues in the GTEx data (40849 domain-contain isoforms from 11141 genes) (GTEx-Consortium, 2020). I find that reference and non-reference domain isotypes are found within expressed transcripts at comparable rates across expression thresholds (Figure 1F), suggesting non-reference domain isotypes are frequently used in human tissues. Accordingly, many non-reference isotype domains were also found when only considering the most expressed (major) transcripts from each gene (Figure S1F-1). Interestingly within the major transcripts, where there are differences in which transcript is the most expressed across tissues, I find 379 (24%) protein domains where the most used domain isotype also changes across tissues (Figure 1G, S1F-1). Similarly, I found 91 (20.7%) domains with isotype switches across a small selection of 9 human cell types profiled at single-cell resolution (Hagemann-Jensen et al., 2020) (Figure 1G). Thus, domain isotypes, created by alternative splicing, are not an exception but the norm and seem to be used in a cell and tissue-specific manner.

### Domain isotypes have distinct 3D structures

Having shown that domain isotypes are prevalent, I next asked if different domain isotypes have different biological functions. Following the “structure is function” axiom (Abbot, 1916), I started with the three-dimensional (3D) structure of the protein domain. I analyzed all human protein domains within the experimentally solved 3D structures in the Protein Data Bank (PDB)(Berman et al., 2000) (Table S3). For each protein domain, I compared the 3D structure of a reference domain isotype to the structure of other reference and non-reference isotypes (as illustrated by arrows in Figure 1D). In other words, I investigated if the 3D structure of protein domain isotypes had evolved through evolution. An interesting example is found in the structure of the histidine phosphatase domain “His_Phos_1”. This domain removes phosphorylations from other proteins via its essential histidine residue (“His_8_”) aided by a few other residues, including “His_108_” (Rigden, 2007). While the “His_Phos_1” protein domains were virtually identical to each other within isotype category (mean RMSD_aligned-backbone_ <1), the reference and truncated isotypes were strikingly different (mean RMSD_aligned-backbone_ >11)(Figure 2A, S2A-1). This difference is exacerbated by the absence of the conserved catalytical residues “His_108_” (Rigden, 2007) from the truncated isotype (shown in blue in the reference isotype domains of Figure 2A, S2A-1). The absence of a conserved catalytical residue naturally implies that the truncated isotype modifies the functionality of the domain (Ma et al., 2005). Importantly this is not just an artifact of the domain detection, as the residue is missing even when considering an additional 50 amino acids downstream of the domain boundary (Figure S2A-2). Encouraged by this example, I summarized the structural differences of domain isotypes across the 952 protein domains analyzed (8741 PDB files, Table S4). I find the 3D structures of different domain isotypes are excessively different (Figure 2B, S2B-1)(Median RMSD_aligned-backbone_ >3x higher than within reference isotype comparison, *P* < 2.14e-102, Wilcoxon tests). Surprisingly specific categories of domain isotypes are associated with larger structural differences that approach what is expected when comparing random domains (shaded area in Figure 2B, Table S5). In summary, the 3D structure of domain isotypes, created through evolution, is surprisingly different, thus challenging the current view of domains as invariable evolutionary building blocks.

**Figure 2:**
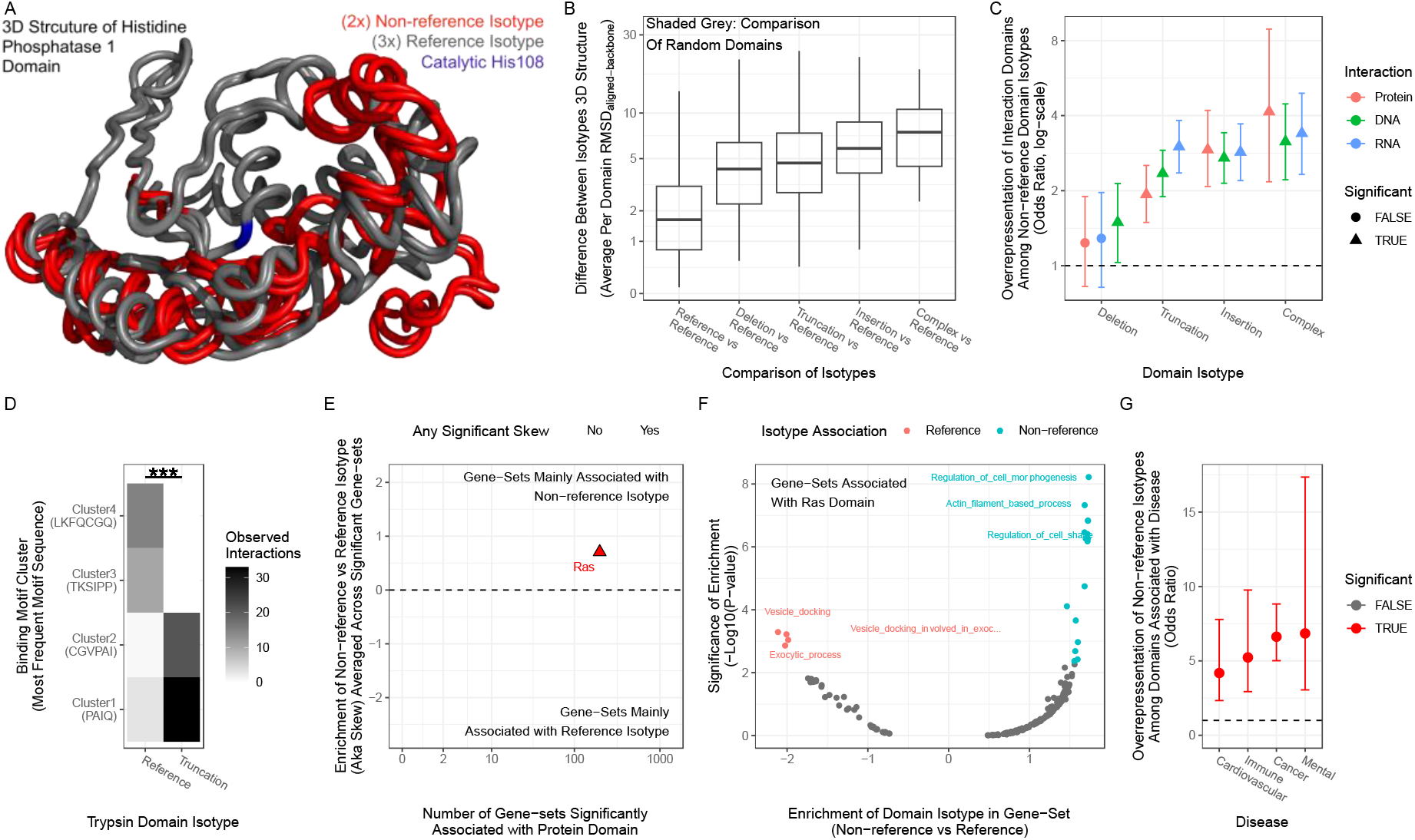
Domain isotypes are functionally important. **A)** Structural alignment of three references (grey) and two truncated (red) isotypes of the His_Phos_1 protein domain. The catalytic residue His108 is indicated in blue. **B)** For each protein domain, the 3D structures of reference isotypes were compared to the structure of all identified isotypes (x-axis). The structural difference was quantified RMSD and averaged per domain (Y-axis) (higher means more different). The grey area indicates the RMSD expected for random domains by showing where the middle 90% of random comparisons fall. **C)** The enrichment (y-axis) of domains involved in interactions (color) for each domain isotype (x-axis). Triangles indicate FDR < 0.05. **D)** The number of interactions (color) found between trypsin domain isotype (x-axis) and clusters of amino acid binding motifs (y-axis) in experimentally solved 3D structures. For each cluster, the most frequent motif is indicated in brackets. **E)** For each protein domain, the number of significantly associated gene sets (x-axis)(FDR < 0.05) as a function of the median skew towards reference or non-reference domain isotype (y-axis). Color indicates if at least one gene set was significantly skewed towards an isotype (FDR < 0.05). **F)** For each-gene set significantly associated with the RAS domain, the skew towards reference or non-reference domain isotype (x-axis) and the associated certainty of the shift (p-value, y-axis). Color indicates significant skew (FDR < 0.05), and top 3 skewed gene-sets in each direction are highlighted. **G)** The enrichment of non-reference domain isotypes (y-axis) among domains associated with various disease classes (x-axis). Color denotes FDR < 0.05.

### Domain isotypes affect biological function

Since interaction between proteins and other macromolecules (e.g., protein, DNA, RNA) requires a specific 3D structure, the structural differences of domain isoforms could modify such interactions (both positively and negatively). In support of this, I find that in Swiss-Prot non-reference protein domain isotypes are highly enriched for protein domains that facilitate interactions with DNA, RNA, and proteins (Mosca et al., 2014) (overall odds ratio per type of interaction > 1.93, p-value < 3.23e-10) (Figure 2C)(Table S6). Indeed, by analyzing the specific amino acid motif that interacts with protein domains in solved 3D structures from DPB (Mosca et al., 2014), I found 10 cases (10.2% of analyzed) where domain isotypes were significantly associated with different clusters of binding motifs (see Methods for details). One example is the trypsin binding domain, where the reference and truncation isotype bind two distinct motif clusters with high specificity (FDR = 8.68e-18, fisher test) (Figure 2D). Thus, protein domain isotypes created through evolution are important for determining the interaction between proteins and other macromolecules, making them particularly interesting from a drug-developmental point of view (Scott et al., 2016).

This is further emphasized since domain isotypes are also created through splicing. Indeed, the GENCODE v38 (Frankish et al., 2020) annotation suggests the human genome contains at least 7735 protein-coding genes (38.71%) that encode protein isoforms, which differ in the presence, absence, or isotype of interaction domains (Figure S2D-1, S2D-2). Thus, protein domain isotypes likely also modulate the function of individual genes through alternative splicing.

The structure is function axiom also suggest that protein domain isotypes could have distinct biological functions. We, therefore, analyzed if individual Swiss-Prot protein domains were overrepresented in the genes annotated to specific pathways or go-terms. I found a significant association between 552 protein domains and 6018 gene sets (FDR < 0.05, Table S7), suggesting the protein domains are critical for the associated pathways. Next, I tested whether the significant gene-sets were preferentially enriched for reference or non-reference domain isotypes (see methods). I found that 77 protein domains had at least one gene set significantly skewed towards an isotype (Figure 2E, Table S5), suggesting that a particular domain isotype was preferred in that pathway. One of these is the Ras protein domain, which is significantly associated with 198 gene-sets, of which 20 were significantly skewed towards a domain isotype (Figure 2F). Interestingly the genes with the reference isotype of the Ras domain seem to be preferentially involved in vesicle formation and release, whereas the genes containing the non-reference isotypes were associated with cell movement (Figure 2F). Thus, domain isotypes might facilitate different biological functions.

Next, I utilized that some protein domains have already been associated with human diseases through overrepresentation of either mutations or naturally occurring disease-related genetic variations (SNPs)(Peterson et al., 2017; Savojardo et al., 2021). I find that non-reference protein domain isotypes are highly enriched for protein domains associated with all 17 disease groups tested (Figure S2G-1), with prominent examples being cardiovascular diseases, mental disorders, immune pathologies, and cancer (all having odds ratio > 4, P-value < 2.33e-07)(Figure 2G). Importantly this suggests that many non-reference domain isotypes are required for homeostasis and, when disrupted, can lead to a wide range of human diseases.

## Discussion

I show that protein domain isotypes are created through both alternative splicing and evolution. These domain isotypes are ubiquitous and modulate or disrupt the functionality of protein domains in a cell, tissue, and disease-specific manner.

Despite the natural dichotomy between alternative splicing and evolution, I here present a joint analysis of both mechanisms to comprehensively examine domain isotypes. Surprisingly these results highlight the synergy of considering both mechanisms since both show domain isotypes are prevalent and functionally important.

However, long-read technology consistently demonstrates that the current databases underrepresent protein diversity (Glinos et al., 2021; Gupta et al., 2018), meaning I probably underestimate the prevalence of domain isotypes. Additionally, naturally occurring genetic variation (SNPs) affect splicing in most human protein-coding genes (GTEx-Consortium, 2020), likely leading to additional domain isotype usage. Jointly this indicates that my results underestimate the true diversity and importance of domain isotypes. Therefore, considerable additional computational and experimental work is needed to characterize the functional role of domain isotypes, both as groups and for individual protein domains.

This article challenges the current perception of protein domains as building blocks with a fixed function. Instead, my findings suggest that protein domain isotypes are via alternative splicing or through evolution used to modulate domain function, thereby increasing the flexibility of protein adaptation. These findings also suggest caution when transferring concussions about a protein domain from one setting to another (both other proteins, cells/tissues, and isoforms) as it is likely that different domain isotypes are used, whereby functional differences are expected. This is especially important when using protein domains for enzyme discovery (Robinson et al., 2021) or drug design (Doğan et al., 2021; Scott et al., 2016).

The widespread use of protein domain isoforms also highlights, along with many other considerations (Barnkob et al., 2022; GTEx-Consortium, 2020; Ji et al., 2020; Vitting-Seerup and Sandelin, 2017), the need to change how we think about molecular biology. Although it is a considerable effort, we must progress from the current “gene-centric” research paradigm to a more nuanced “isoform-centric” paradigm.

## Methods

### Data and statistical analysis

All analysis was done in R 4.0.4 unless explicitly stated. Significance is in all analyses defined as Benjamini-Hochberg (BH)(also known as False Discovery Rate, FDR (Benjamini and Hochberg, 1995)) corrected p-values smaller than 0.05.

### Domain Analysis

All domains analyzed in this article were identified by running Pfam’s (Mistry et al., 2020) pfam_scan.pl tool on the corresponding amino acid sequences only using the manually curated pfamA database(Mistry et al., 2020) and default parameters. All results in this article are only based on protein domains defined by the “type” column of the pfam_scan output (meaning no families, repeats, etc., were considered).

### Calculating Domain Variation Sizes

To analyze domain sizes, I use the hmm coordinates, alignment coordinates, and the reference domain length, all available from the output of pfam_scan.pl. Truncation sizes for each end of the domain are calculated as respectively hmm_start -1 and hmm_end -hmm_length, and the largest value is used as the truncation size. The truncation fraction is calculated by dividing the truncation size by the hmm_length, defined as hmm_end – hmm_start + 1.

Indel sizes are extracted by comparing the length of the aligned sequence and the aligned reference model. Specifically, the length of the aligned reference model is calculated as hmm_end – hmm_start + 1, and the length of the aligned sequence is calculated as aligned_end – aligned_start +1. The indel size is calculated by subtracting the aligned sequence length from the aligned model length, and the indel fraction is calculated by dividing by the aligned model length.

The overall domain variation is defined as the largest absolute fraction when comparing the truncation fraction and indel fraction. For the cluster analysis, I used the Modes() function from the LaplacesDemon R package modifying the “min.size” argument as indicated in the figure (0.1-0.3). The Modes function finds the number of local maxima in the density distribution of the variables while ensuring a mode corresponds to a least the fraction of observations indicated by the “min.size” argument.

### pfamAnalyzeR

I here implemented and used pfamAnalyzeR v. 0.1.0 (available at Bioconductor (Huber et al., 2015) https://bioconductor.org/packages/pfamAnalyzeR/), which currently has three functionalities: 1) It reads the fixed-with-file format produced by pfam_scan.pl or the Pfam webserver into R taking know format problems and data versions into account. 2) Domain data is augmented with the domain truncation/indel calculations described above. 3) Each domain is assigned one of 5 mutually exclusive isotype categories. First, a domain is defined as having a truncation if the truncation fraction >= 0.1 (Figure S1D-1). Secondly, a domain is defined as having an indel if the absolute indel fraction is >= 0.1 (Figure S1D-1). Then the 5 mutually exclusive categories are defined as:

1. “Reference”: No truncation and no indel
2. “Complex”: Both truncation and indel
3. “Truncation” : Truncation but no indel
4. “Insertion”: No truncation but indel and indel fraction > 0
5. “Deletion”: No truncation but indel and indel fraction < 0

These classes are further simplified to “reference” (#1 from above) and “non-reference” (#2-#5 from above).

### TCGA Analysis

The TCGA isoform switches were directly obtained from the supplementary material of Vitting-Seerup *et al*. (Vitting-Seerup and Sandelin, 2017). I extended IsoformSwitchAnalyzeR (Vitting-Seerup and Sandelin, 2019) to incorporate results from pfamAnalyzeR and to compare the simplified isotype classification (reference vs. non-reference) as a new functional consequence (available in IsoformSwitchAnalyzeR v1.17.06). The number of genes with isoform switches was extracted with the extractConsequenceSummary() function. The ABI1 visualization was created with the switchPlot() and switchPlotTranscript() functions. Data is available in Table S1.

### Species Analysis

The amino acid fasta file for the manual curated SwissProt subset of UniProt (UniProt-Consortium et al., 2020) was downloaded from UniProt’s website, downloading only canonical sequences (one representative sequence per gene). The human file was downloaded on 11^th^ October 2021. The Saccharomyces Cerevisiae, Mus Musculus, and Drosophila Melanogaster files were downloaded on 12^th^ May 2022. All files were analyzed with pfam_scan and pfamAnalyzeR as described above. Domains identified, along with their isotype, are available in Table S2.

### Tissue Expression analysis

The GTEx transcript TPM expression for all 55 human tissues was downloaded from gtexportal.org using v8 data (GTEx-Consortium, 2020). For each tissue, the median transcript expression was calculated. Since the GTEx v8 data is quantified via GENCODE (Frankish et al., 2020) v23 annotation, I obtained the corresponding amino acid from gencodegenes.org and analyzed it with pfam_scan and pfamAnalyzeR as described above (Table S8). The major transcript is defined as the most expressed protein-coding transcript from a given gene. For Figure 1G, I analyzed genes with two different transcripts defined as major in at least one tissue. Domain switches were then identified as those with differences in which domain isotypes were encoded in the transcripts.

### Cell Type Expression analysis

The differential transcript usage analysis across 9 human immune cell types identified from full-length single-cell data was downloaded from the supplementary data Hagemann-Jensen *et al*. (Hagemann-Jensen et al., 2020) (the “noHEK” analysis). Transcripts with usage changes were extracted by filtering for adjusted p-values < 0.05. For each significant transcript, I identified the cell type where it was most expressed. For Figure 1G, I analyzed genes where two different transcripts are defined as major in two different cell types. Domain switches were then identified by analyzing the isotypes of the domains encoded in the corresponding GENCODE v23 transcripts.

### Structural analysis

I used PDB’s (Berman et al., 2000) batch_download.sh, script to download all human pdb files 13^th^ of October 2021. From these pdb files, I extracted the amino acid sequence of the first chain (not considering X) using the read.pdb() function from the bio3d R package(Grant et al., 2020). The resulting fasta file was analyzed with pfam_scan and pfamAnalyzeR as described above. Domains identified, along with their isotype, are available in Table S3.

It was not computationally feasible to analyze all possible within-domain combinations. Instead, for each domain (hmm_name), I made all pairwise comparisons of max 9 randomly selected reference isotypes and max randomly selected 21 non-reference isotypes. The selected reference isotypes were also compared to each other. To estimate what is expected by chance, I randomly selected one structure for 150 different domains, all classified as the reference isotype, and extracted all possible pairwise comparisons.

All structural comparisons were made using the bio3d R package(Grant et al., 2020) v 2.4-2. The pdb files were loaded into R using the read.pdb(). The correct chain and the subset of the structure containing the domain were extracted using the atom.select() function with the Pfam domain alignment coordinates. The structures were compared using the struct.aln() function without trimming optimization (max.cycles=0). This function works by doing a sequence alignment and afterward using the aligned sub-sequence to rotate and transpose the 3D structures. From this structural alignment, the function calculates the backbone RMSD, and the function was manually extended to calculate the backbone TM-scores as defined in equations 1 and 5 in Zhang *et al*. (Zhang and Skolnick, 2004). The sequence alignment was run with the following parameters: gapOpening = 40, gapExtension = 4, substitutionMatrix = BLOSUM50 (from the Biostrings package) to encourage an alignment better reflecting the expected input where large parts were missing from ends (truncation) or in the middle of the sequence (insertion/deletion).

I only analyzed comparisons where both of the aligned sequences were > 20 amino acids and where the length of the non-reference aligned sub-sequence corresponded to >80% of the original sequence length (n= 48369)(Table S4-5). To obtain the average structural differences I for each domain (hmm_name) and isotype extracted the mean RMSD and TM-score for the cases where at least 3 structural comparisons passed the filtering described above. Average TM-scores and RMSD values are highly negatively correlated (spearman’s rho: 0.891, p-value < 2.22e-16). The area of random RMSD values was defined as the 5-95 percentile of all 11026 random comparisons.

For the His_Phos_1 example 3 reference isotype structures (PDB ID: 1K6M(Lee et al., 2003), 5HTK(Crochet et al., 2017), 6HVH(Boutard et al., 2019)) and 2 truncation isotype structures (PDB ID: 1YFK(Wang et al., 2005), 2A9J(Wang et al., 2006)) were manually selected. Note that these are from 5 different genes, and “only” 5 structures were chosen because this was the maximal number I could get the multiple alignment described below to work with. The pdb files and domain subsets were handled as described above. All 5 domain structures were aligned with the pdbaln() function using the abovementioned parameters. This function works the same as struct.aln(), except it uses multiple alignment to align 2+ sequences. Open-Source PyMOL (https://github.com/schrodinger/pymol-open-source) was used to visualize the aligned structures by saving them with pymol.pdbs() and afterward opening it in PyMOL. The catalytic residues were manually annotated by comparison to Figure 3 in Rigden *et al*. (Rigden, 2007) using the 1K6M structure as reference since it is included in both analyses.

### Domain Interaction Analysis

Protein domains interacting in solved 3D structures were downloaded from the newest version (2020_01) of the 3DID database(Mosca et al., 2014) (https://3did.irbbarcelona.org). DNA and RNA binding domains are defined as domains returned by searching the Pfam website (https://pfam.xfam.org/) for, respectively, “DNA binding” and “RNA binding” keywords. See Table S6.

For the interaction analysis, I only analyzed domains found in human SwissProt. For each interaction type, I used Fisher’s exact test to test the overlap between each domain isotype and the domains associated with the interaction type.

### Binding Motif Analysis

For the domain-motif interaction (DMI), I downloaded the 3DID DMI (Mosca et al., 2014) from https://3did.irbbarcelona.org. The DMI data was reduced to the interaction containing the human pdb files and chains I used for the structural analysis described above. The data was further reduced to domains uniquely overlapping (in protein coordinates) of the same type (hmm_name), and the domain isotype was transferred from the pdb data to the DMI data.

For each domain (hmm_name), I clustered motifs using the distance matrix produced by stringdistmatrix() from the stringDist R package(Loo, 2014) as input to standard hierarchical clustering with hclust(). Clusters were defined using the cutree() function specifying h = 4. The interpretation is that clusters are sets of motifs where >4 modifications (deletions, insertions, substitutions, and transpositions) must be made to make the motifs similar to any motif outside the cluster. Considering that the median motif consists of 6 amino acids, this results in very different clusters.

To identify domains where isotypes were associated with distinct motifs clusters, I only considered motif clusters where for each domain (hmm_name), at least one domain isotype was identified at least 3 times. A motif cluster was defined as isotype-specific if 80% of motifs within the cluster interacted with either reference or non-reference domain isotypes. Of the 98 domains (hmm_name) with at least two motif clusters, I found 10 where reference and non-reference isotypes were specific to distinct motif clusters. A Fisher test was used to test the statistical significance of the Trypsin example using the 4 × 2 contingency table of observed interaction counts where “4” indicate the 4 motif clusters and “2” is reference vs. non-reference groups.

### Differences in presence, absence, or isotype of interaction domains

To determine changes in the presence, absence, or isotype of interaction domains, I used Pfam to identify protein domains (as described above) in the GENCODE v38 (Frankish et al., 2020) annotation. I then extracted genes containing the interaction protein domains (as defined above). I then counted how many genes had isoforms both with and without a protein domain (n = 6638) and how many genes had protein domains (same hmm_name) that overlapped in genomic coordinates but were classified as different domain isotypes by pfamAnalyzeR (as described above).

### Gene-set association

I used enrichment analysis to associate specific domains (given by their hmm_name) with biological functions. First, I extracted gene sets from MSigDB (Liberzon et al., 2015, 2011) explicitly using the following collections: 1) Hallmarks, C6 oncogenic, GO:BP Biological Processes part of gene ontology, KEGG pathways, Wikipathways, and CPG collection containing gene sets describing chemical and genetic perturbations. From these, only gene sets with at least 10 and, at most, 1000 gene symbols were used. The domains identified in SwissProt were reduced to those found in at least 10 genes. Both gene sets and domains were reduced to genes found in both datasets and gene sets with <10 genes were again removed. For each domain (hmm_name), I use the Fisher.test() function with alternative= “greater” to evaluate the enrichment of the genes in which the domain was found among the genes in a gene set. For each domain, P-values were corrected for multiple FDR testing, and significant associations were defined as FDR < 0.05.

For each gene set significantly associated with a domain, I tested if the same gene set was also significantly associated with either the reference or the non-reference isotype of that domain as follows. For each gene and domain (hmm_name), I calculated the average overall domain variation (see Domain Analysis section for that definition), where larger values mean larger deviation from the reference isotype. For each domain, I then extracted the subset of genes with that domain and ranked them according to the average overall domain variation in decreasing order. I then used fgsea (Korotkevich et al., 2021) to test if the significant gene set was overrepresented at the top or bottom of the list (skew). The decreasing ranking means that positive NES scores indicate associations with non-reference isotypes, and negative NES scores indicate associations with reference isotypes. Gene sets were only tested if there was at least one gene in the ranked list that was not in the gene set. For each domain, P-values were corrected for multiple testing using FDR, and significant associations were defined as FDR < 0.05. The gene sets associated with each domain (and isotype skew) are available in Table S7.

### Disease association

I obtained the Pfam domain associated with diseases from supplementary table 4 of Savojardo *et al*. (Savojardo et al., 2021) by filtering for associations > 0. Protein domains associated with cancers (oncodomains) were obtained from the supplementary data of Peterson *et al*. (Peterson et al., 2017) and added to the list of disease-associated domains. The disease-associated domains were reduced to those found in the human SwissProt data. For each disease group, I then used a Fisher’s exact test to test the overlap between each isotype and the domain associated with the disease group. Similarly, another test was performed where all non-reference isotypes were jointly analyzed. P-values were corrected for multiple testing using FDR, and significant associations were defined as FDR < 0.05.

## Acknowledgments

Albin Sandlin for the encouraging discussion of the initial idea. For insightful feedback on manuscript drafts, Elena Papaleo, Lars Rønn Olsen, previous reviewers, and Bo Porse.

