## Supplementary figures for "Most protein domains exist as variants with distinct functions across cells, tissues, and diseases"

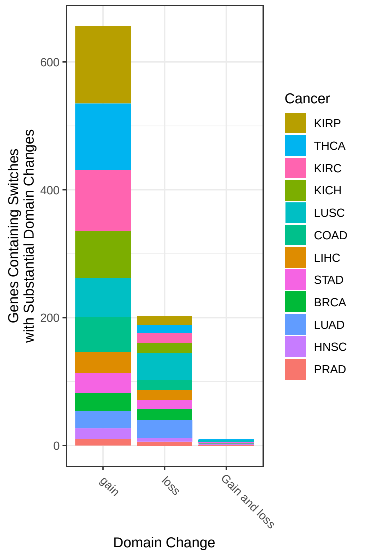


**Figure S1A-1.** For each cancer type (color), the number of genes with partial gain/loss of protein domains (y-axis). Genes are divided based on whether the isoform switch resulted in a gain, a loss, or both gain/loss of a parts of the protein domain (x-axis). BRCA: Breast invasive carcinoma, COAD: Colon adenocarcinoma, HNSC: Head and Neck squamous cell carcinoma, KICH: Kidney Chromophobe, KIRC: Kidney renal clear cell carcinoma, KIRP: Kidney renal papillary cell carcinoma, LIHC: Liver hepatocellular carcinoma, LUAD: Lung adenocarcinoma, LUSC: Lung squamous cell carcinoma, PRAD: Prostate adenocarcinoma, STAD: Stomach adenocarcinoma, THCA: Thyroid carcinoma.

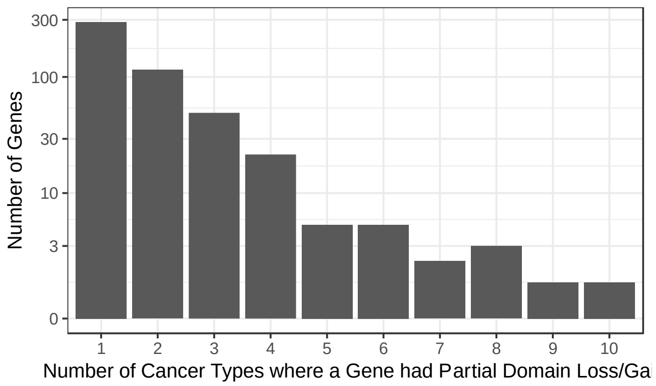


**Figure 1A-2. 2)** The number of cancers in which each gene from figure Figure S1A-1 was identified.

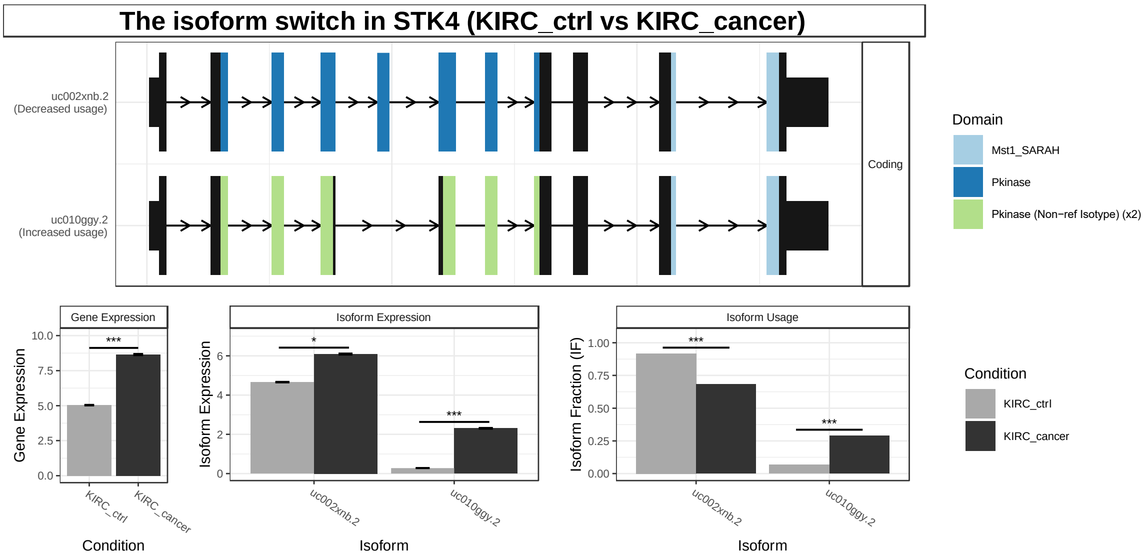


**Figure S1A-3.** The switchPlot of STK4 in Kidney Renal Clear Cell Carcinoma (KIRC). Top sub-plots as described in Figure 1A of main text. Bottom row of sub-plots shows the conditional gene expression (left), transcript expression (midt) and isoform fraction (right). Isoform fractions indicate how large a fraction each isoform contribute to the overall gene expression. *** denotes FDR corrected p-values < 0.001. * denotes FDR corrected p-values < 0.05

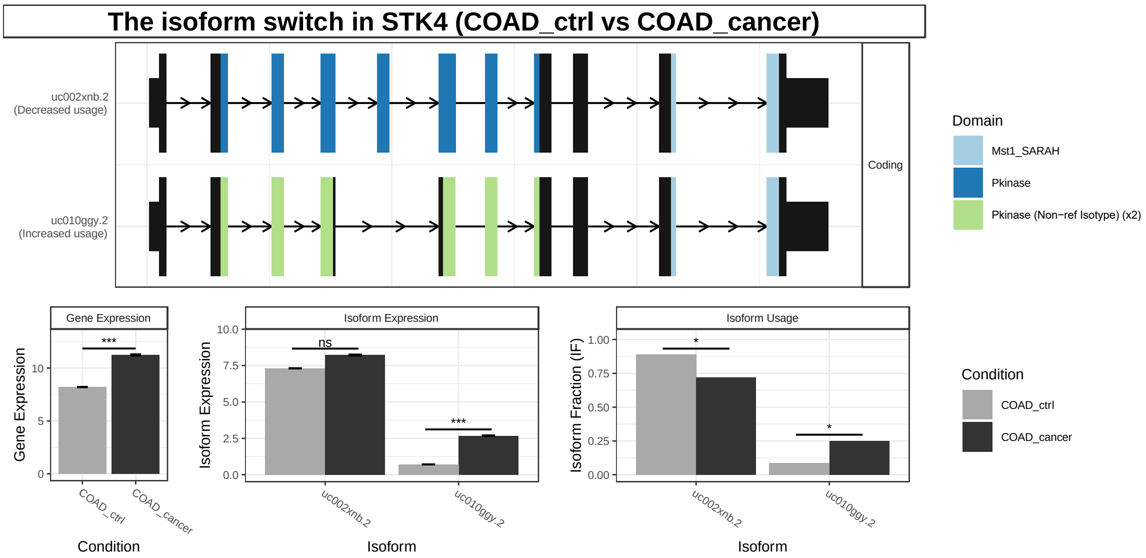


**Figure S1A-4.** The switchPlot of STK4 in Colon adenocarcinoma (COAD) as described in figure S1A-3.

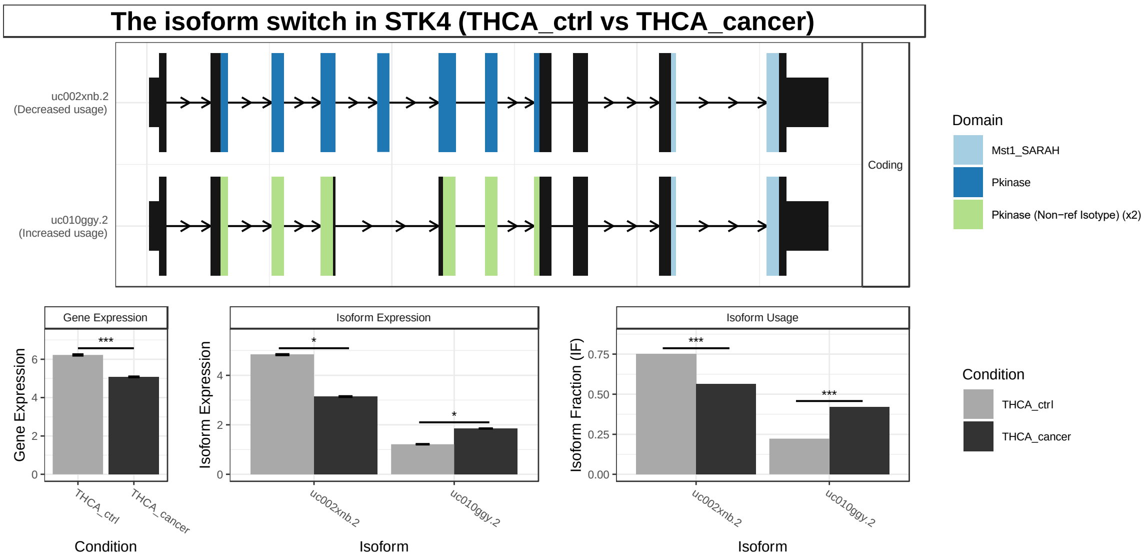


**Figure S1A-5.** The switchPlot of STK4 in Thyroid carcinoma (THCA) as described in figure S1A-3.

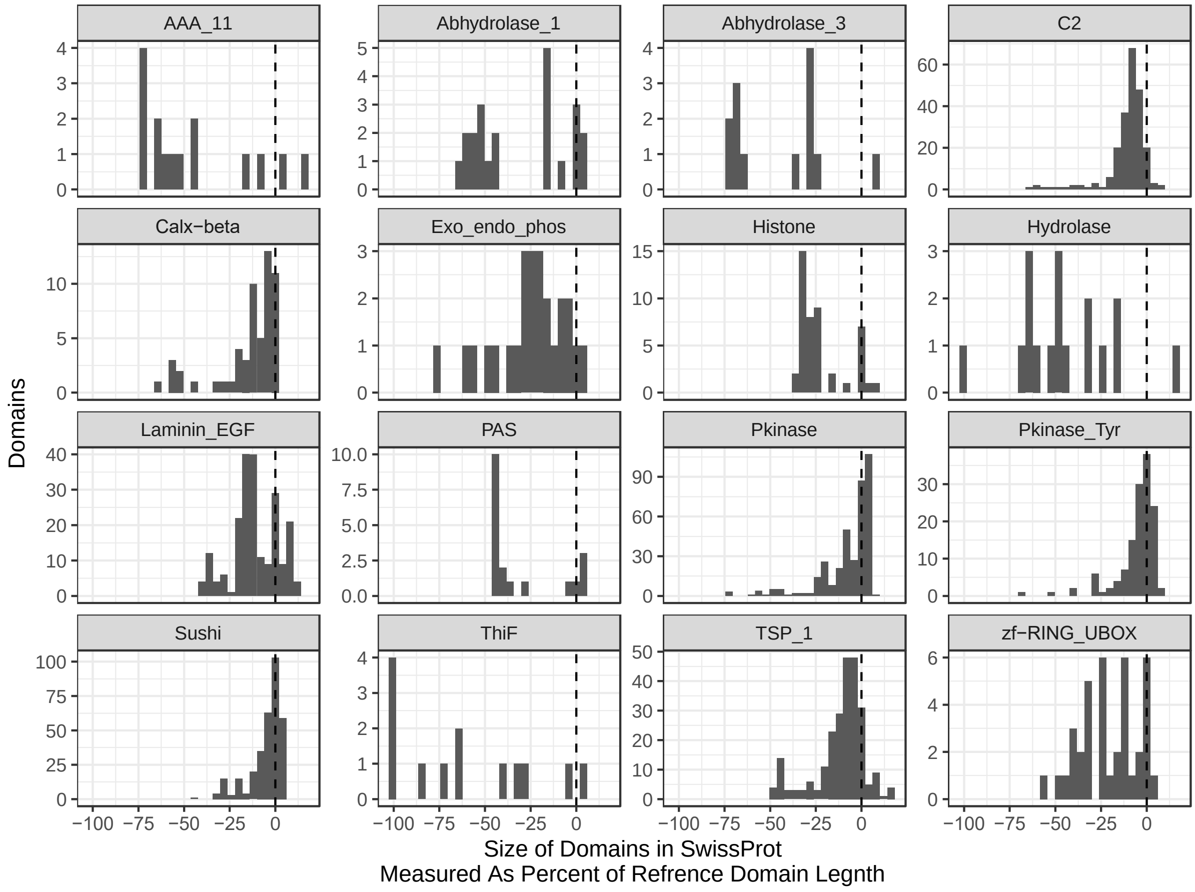


**Figure S1C-1.** The domain length differences of the 16 protein that most frequently have large domain differences (more extreme than 25% of the reference length). The difference is measured as a fraction of the reference length with the dashed line indicating no difference.

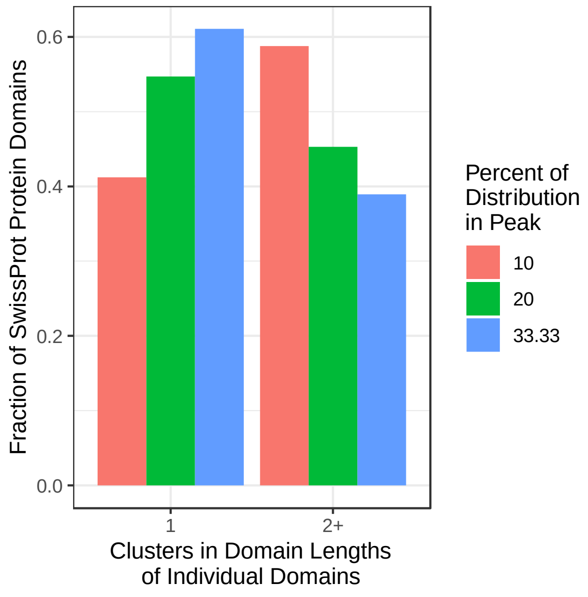

**Figure S1C-2.** The number of clusters found in the length distribution of each protein domain identified in SwissProt. Color indicate the minimum percent of domain lengths in each cluster.

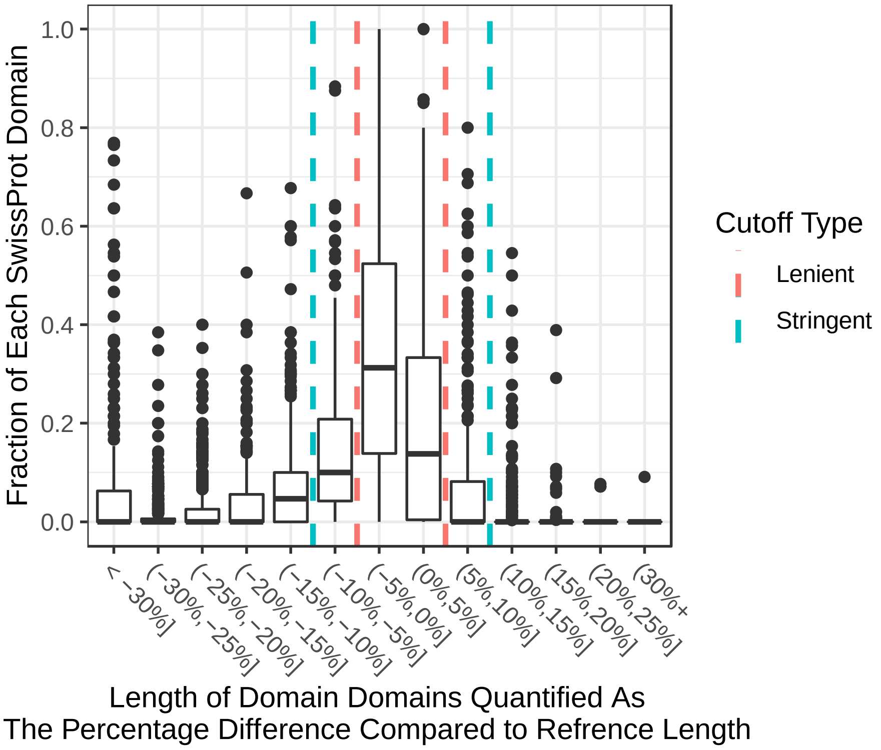


**Figure S1D-1.** The frequency (y-axis) of the difference between observed protein domain length and the corresponding reference model measured as a fraction of the reference length. The length differences across all domains identified in SwissProt and each bin is shown as a boxplot. Length differences capped at +/- 30% for visual purposes. Stringent and lenient cutoffs for defining isotypes are shown.

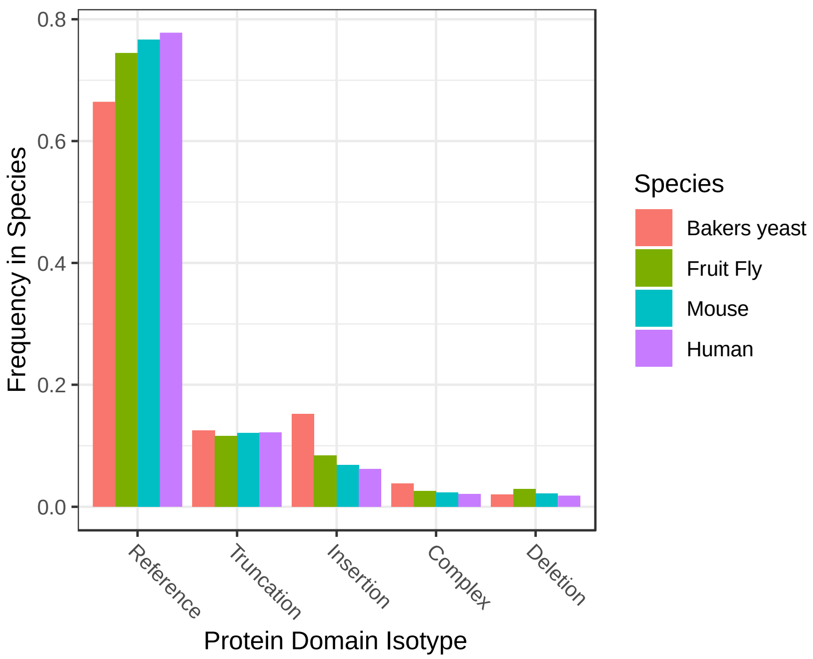


**Figure S1E-1.** The frequency (y-axis) of each protein domain isotype (x-axis) in SwissProt proteins for different species (color).

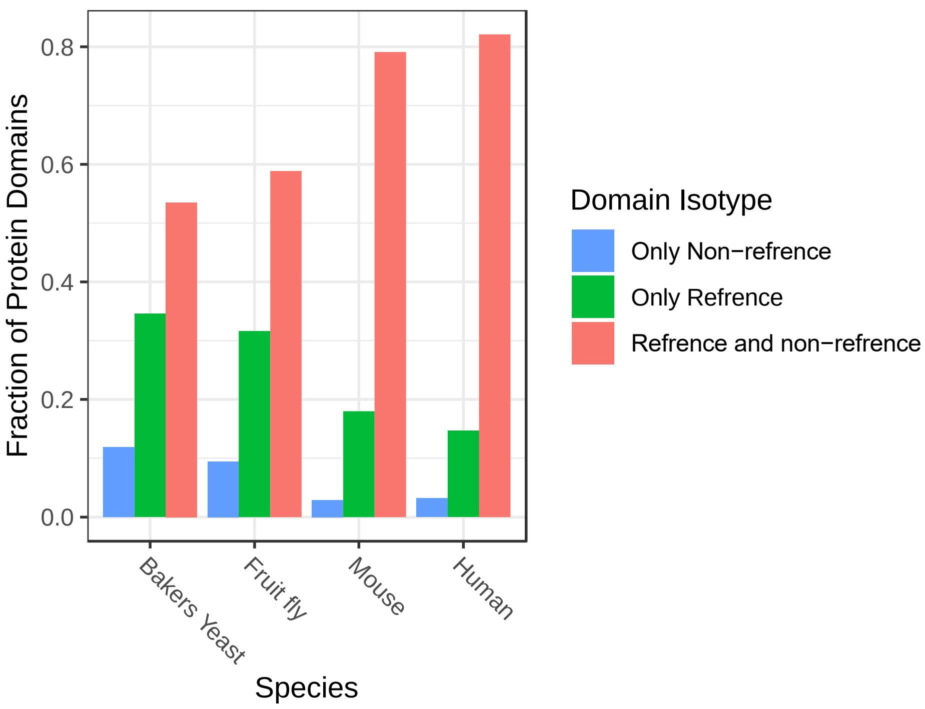


**Figure S1E-2.** The fraction of protein domains that exist as reference and non-reference isotypes in SwissProt of various species when only considering proteins annotated as "domain containing" in SwissProt.

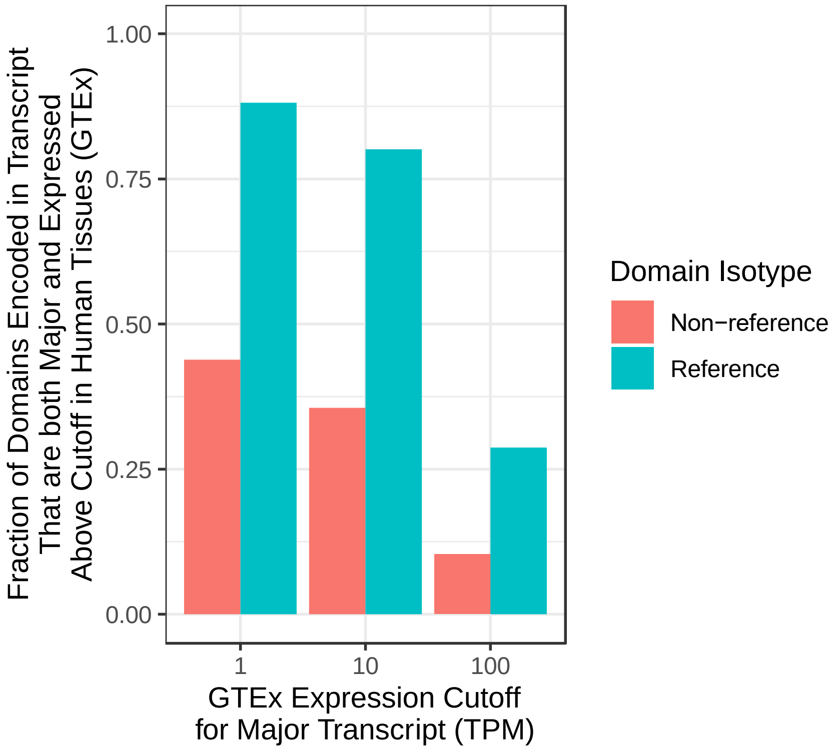


**Figure S1F-1.** The fraction (Y-axis) of domain isotypes (color) contained in a major transcript also expressed above a Transcript Per Million (TPM) cutoff (x-axis) in at least one human tissue. A major transcript is defined as the transcript that is most expressed from its gene in at least one tissue type.

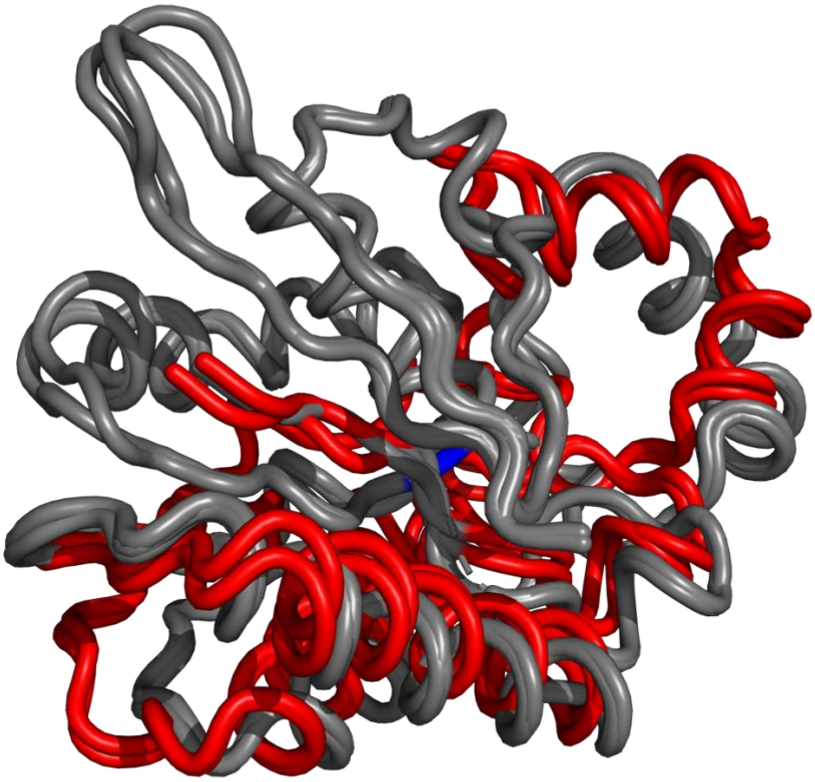

**Figure S2A-1.** Structural alignment of three references (grey) and two truncated (red) isotypes of the His_Phos_1 protein domain. The catalytic residue His108 is indicated in blue. The same as in the main figure just shown from another angle.

```
                  1        .         .         .         .         .         .         70
1K6M_A (ref)      IYLCRHGESELNIRGRIGG--DSGLSVRGKQYAYALANFIQSQGISSLKVFTSRMKRTIQTAEALGVPYE
5HTK_A (ref)      IYLCR-GESEFNLLGKIGG--DSGLSVRGKQFAQALRKFLEEQEITDLKVWTSQLKRTIQTAESLGVPYE
6HVH_A (ref)      IYLCRHGENEHNLQGRIGG--DSGLSSRGKKFASALSKFVEEQNLKDLRVWTSQLKSTIQTAEALRLPYE
1YFK_A (non-ref)  LVLIRHGESAWNLENRFSGWYDADLSPAGHEEAKRGGQALRDAGYEFDICFTSVQKRAIRTLWTVLDAID
2A9J_A (non-ref)  LIMLRHGEGAWNKENRFCSWVDQKLNSEGMEEARNCGKQLKALNFEFDLVFTSVLNRSIHTAWLILEELG
                  ^ ^ * **   *   ^     *  *   *   *      ^          ^**     * *   ^
                  1        .         .         .         .         .         .         70

                  71        .         .         .         .         .         .         140
1K6M_A (ref)      QFKALNEIDAGVCEEMTYEEIQEHYPEEFALRDQDKYRYRYPKGESYEDLVQRLEPVIMELERQENVLVI
5HTK_A (ref)      QWKILNEIDAGVCEEMTYAEIEKRYPEEFALRDQEKYLYRYPGGESYQDLVQRLEPVIMELERQGNVLVI
6HVH_A (ref)      QWKALNEIDAGVCEELTYEEIRDTYPEEYALREQDKYYYRYPTGESYQDLVQRLEPVIMELERQENVLVI
1YFK_A (non-ref)  QMWLPVVRTWRLNERHYGGLTGLNKAETAAKHGEAQVKIWRRSYDVPPPPMEPDHPFYSNISKDRRYA--
2A9J_A (non-ref)  QEWVPVESSWRLNERHYGALIGLNREQMALNHGEEQVRLWRRSYNVTPPPIEESHPYYQEIYNDRRYKVC
                  *          ^ *                 ^                  ^    *    ^
                  71        .         .         .         .         .         .         140

                  141        .         .         .         .         .         .         210
1K6M_A (ref)      CHQAVMRCLLAYFLDKSSEELPYLKCPLHTVLKLTPVAYGCKVESIYLNVEAVNTHREKPENVDITREPE
5HTK_A (ref)      SHQAVMRCLLAYFLDKGADELPYLRCPLHTIFKLTPVAYGCKVETIKLNVEAVNTHRDKPTNNFPK----
6HVH_A (ref)      CHQAVLRCLLAYFLDKSAEEMPYLKCPLHTVLKLTPVAYGCRVESIYLNVESVCTHRERSE---------
1YFK_A (non-ref)  --------------DLTEDQLPSCESLKDTIARALPFWNEEIVPQIKEGKRVLIAAHGN-----------
2A9J_A (non-ref)  --------------DVPLDQLPRSESLKDVLERLLPYWNERIAPEVLRGKTILISA--------------
                       *   ^ ^*        ^ ^  *         ^      ^
                  141        .         .         .         .         .         .         210

                  211        220
1K6M_A (ref)      EALDTVPAHY
5HTK_A (ref)      ----------
6HVH_A (ref)      ----------
1YFK_A (non-ref)  ----------
2A9J_A (non-ref)  ----------

                  211        220
```

**Figure S2A-2**. Sequence alignment comparing the structure of 5 His_Phos_1 domains from distinct structures (and genes). The PDB id and chain are given as sequence ids to the left. Domain isotype is given in brackets. The core catalytical residues are in bold, enlarged, and colored red when conserved between isotypes, and blue when missing from the non-reference domain isotype. The grey sequence are the residues aligned when 50 amino acids downstream of the domain boundary are included (note that for the reference isotype this exceeds the end of the protein hence fewer amino acids can be shown in grey).

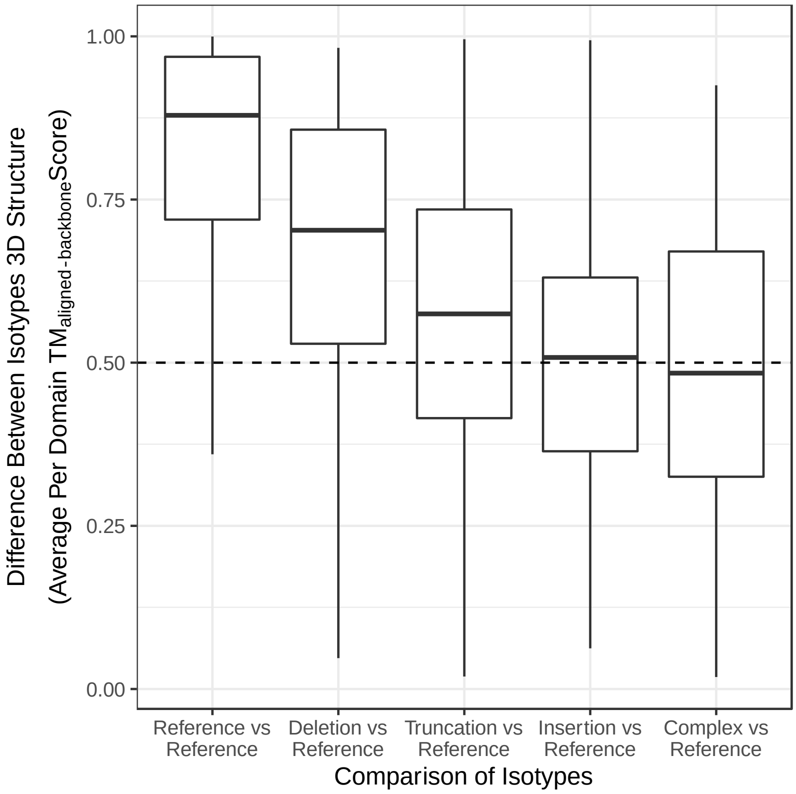


**Figure S2B-1.** For each protein domain, the 3D structures of reference isotypes were compared to the structure of all analyzed isotypes (x-axis). The structural difference was quantified using TM-scores and averaged per domain (Y-axis) (higher scores mean more similar). The dashed line indicates what is expected from structures in approximately the same fold.

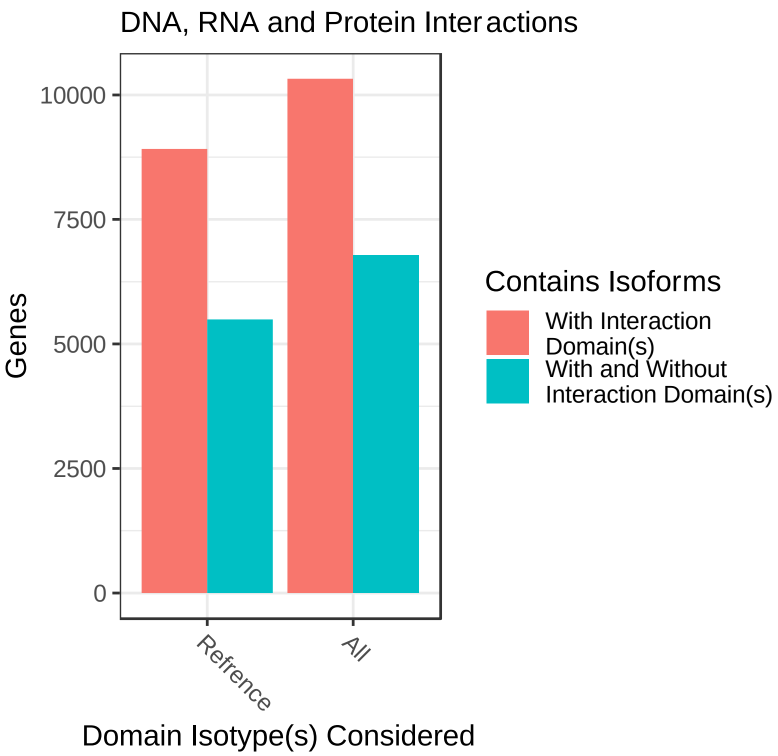


**Figure S2D-1.** The number of GENCODE genes (y-axis) that has isoforms encoding an interaction domain (red) or isoforms that both encode and does not encode an interaction domain (blue) depending on whether only "reference" isotypes are counted or all isotypes are counted (x-axis).

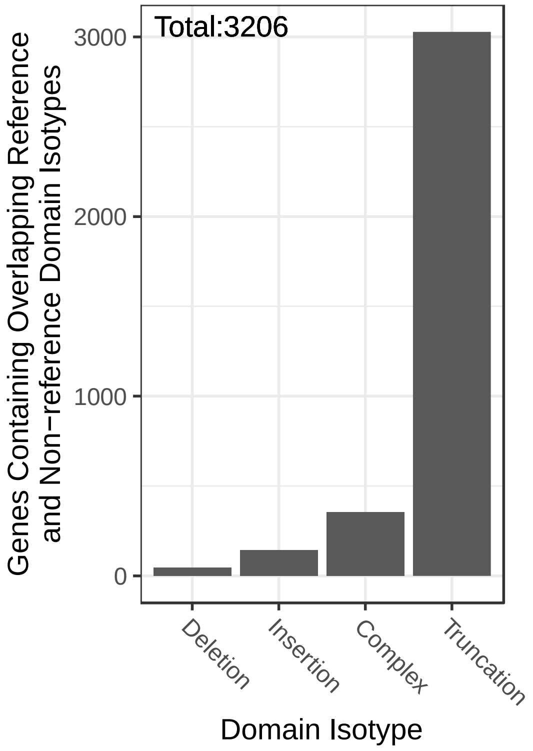


**Figure S2D-2.** For each isotype (x-axis), the number of GENCODE genes that encodes reference and non-reference domain isotypes from the same genomic region (encoded in different isoforms)(Y-axis). The total number of genes encoding at least one such pair is indicated in the top left corner.

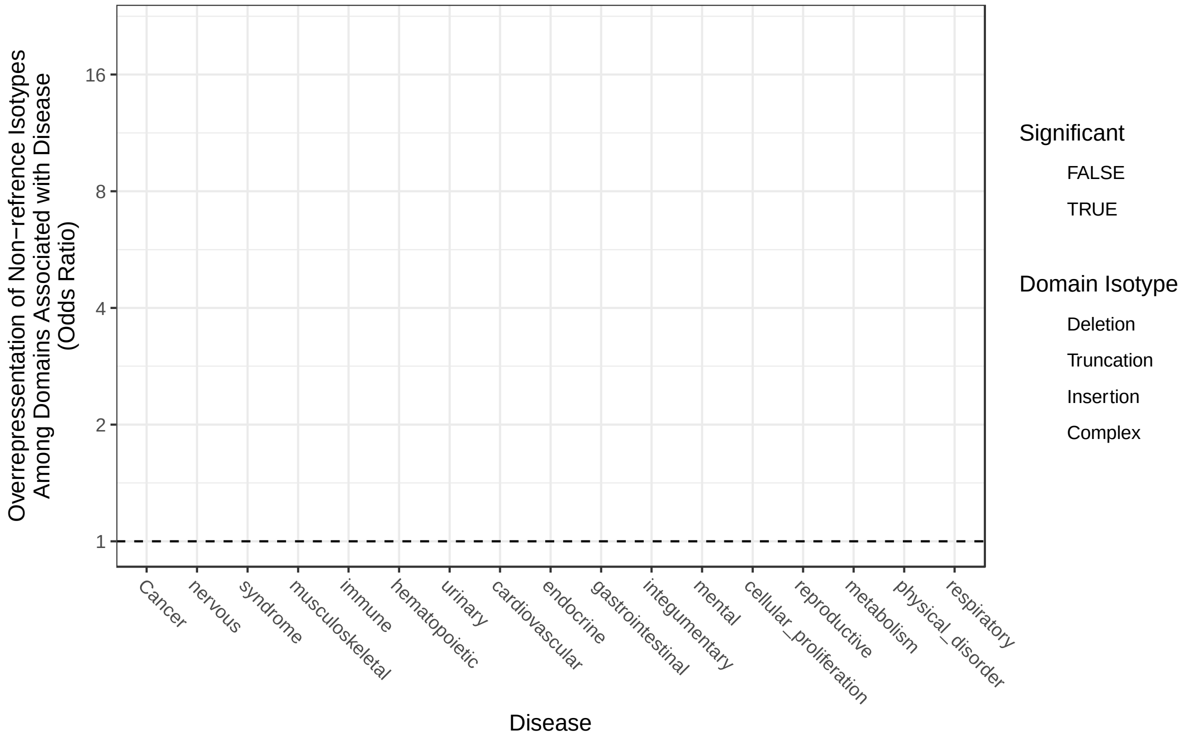


**Figure S2G-1. 1)** The enrichment (y-axis) of non-reference domain isotypes (color) among domains associated with various disease classes (x-axis). Shape denote FDR < 0.05

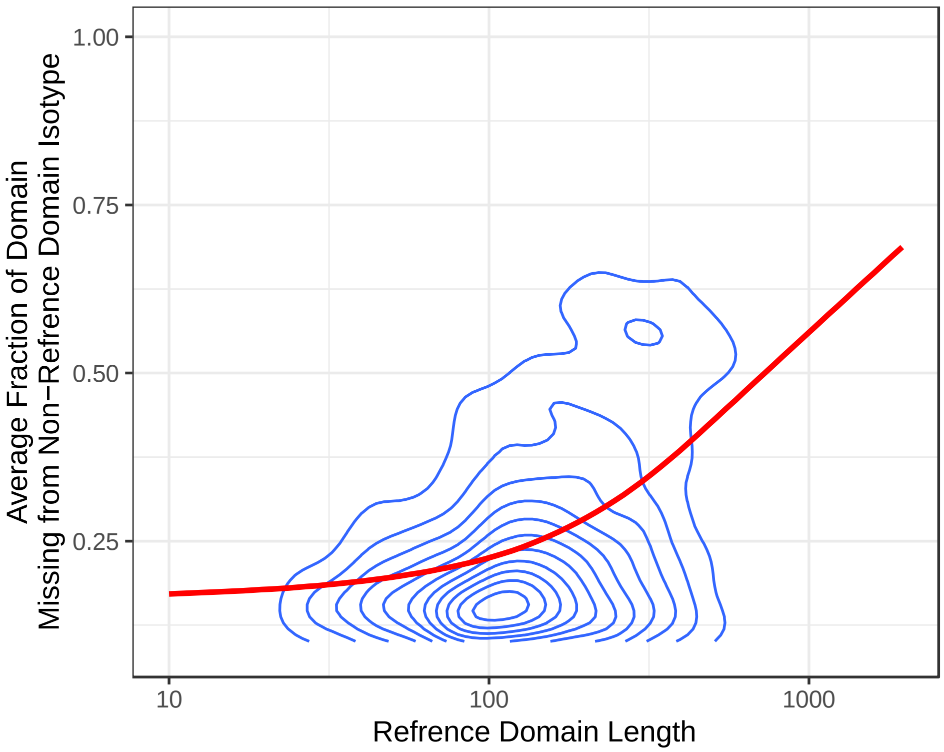


**Figure S3.** The average domain length difference of non-reference domain isotypes (Y-axis) as a function of reference domain lengths (x-axis). The domain length difference is quantified as a fraction of the reference domain length.
